# gene2fish: a multi-gene spatial expression browser for in situ hybridization data across zebrafish development

**DOI:** 10.64898/2026.10.16.759755

**Authors:** Vera Janssen, Wellington D. F. Rutes, Leandro Lima, Kyle Awayan, Benjamin Iovino, Rafael Gómez-Sjöberg, Loïc A. Royer

## Abstract

Single-cell RNA-sequencing (scRNA-seq) atlases of zebrafish development have catalogued marker genes for hundreds of cell types, but transcriptomic data alone cannot reveal their expression patterns in the embryo. The systematically annotated Thisse high-throughput whole-mount in situ hybridization (ISH) collection at ZFIN provides spatial expression data across twelve developmental stages, yet the ZFIN web interface is organized around individual genes, making side-by-side cross-gene comparison laborious. We present gene2fish, a web application that displays ISH images from the Thisse dataset, linked directly to their ZFIN records, in a gene × developmental-stage grid. With gene2fish, researchers can quickly explore and validate the spatial expression of marker gene lists, and find the genes annotated to specific anatomical structures across developmental time. The application is part of Zebrahub and is freely available at https://zebrahub.sf.czbiohub.org/gene2fish.

## Introduction

Zebrafish (*Danio rerio*) is one of the most extensively profiled vertebrates at single-cell resolution. A succession of whole-embryo single-cell RNA-sequencing (scRNA-seq) studies has produced high-resolution transcriptomic maps spanning virtually all of embryogenesis and early larval stages^1–10^. Together, these resources have assigned transcriptomic identities to hundreds of zebrafish cell types and produced large, detailed catalogues of cell-type marker genes. A fundamental limitation of scRNA-seq, however, is the loss of spatial context: dissociation destroys the positional information that defines tissue architecture. Marker genes identified computationally therefore need to be checked against spatial expression data, to confirm cell-type identity, delineate tissue boundaries and generate hypotheses about tissue dynamics and cellular interactions. Spatially resolved transcriptomic approaches, from RNA tomography^11^ and Stereo-seq^12^ to whole-embryo MERFISH^13^ and expansion sequencing^14^, can profile hundreds of genes up to the whole transcriptome and give a broad overview of spatial gene expression. However, these approaches require specialized instrumentation, substantial experimental time and computational resources, and are therefore not yet routine tools for the rapid exploration and validation of marker gene lists. Especially at later developmental stages, spatial gene expression datasets are scarce. Therefore, marker genes are still cross-checked against published RNA in situ hybridization expression patterns.

Currently the largest systematic spatial expression data known for zebrafish is the Thisse *et al*. high-throughput whole-mount ISH collection which is hosted at ZFIN^15–20^. It is an unusually accessible resource covering thousands of genes from blastula to early larval stages. Despite this breadth, ZFIN’s web interface is organized around individual genes and publication figures: although its expression search can filter annotations by anatomy term and stage, there is no facility for viewing the images of multiple genes side by side along a shared developmental axis, or for browsing the images of all genes annotated to one anatomical structure in a single view. Browsing the data one gene at a time therefore limits its value and often restricts its use to zebrafish specialists rather than researchers from other disciplines and the general public. We developed gene2fish to make the gene expression patterns in these in situ hybridization data accessible for visual exploration.

## Results

### gene2fish application

We introduce gene2fish, a web-based tool on the Zebrahub portal that displays RNA in situ hybridization data as a gene-by-stage grid (Fig. 1). gene2fish hosts expression data of whole-mount zebrafish embryos for 8,263 genes across 53,595 images from the Thisse records, with 1–144 images per gene (median 1) spanning sphere stage to larval day 5 (Supplementary Fig. 1A–B). Coverage is concentrated between the 1–4 somite and High-pec stages; germ-ring, shield and pec-fin stages are represented by only one or two genes each, and 4,909 images are annotated to an unspecified stage (Supplementary Fig. 1B); 3,850 genes have at least one stage-resolved image. Developmental stages follow the standard zebrafish staging series^21^.

**Fig. 1:**
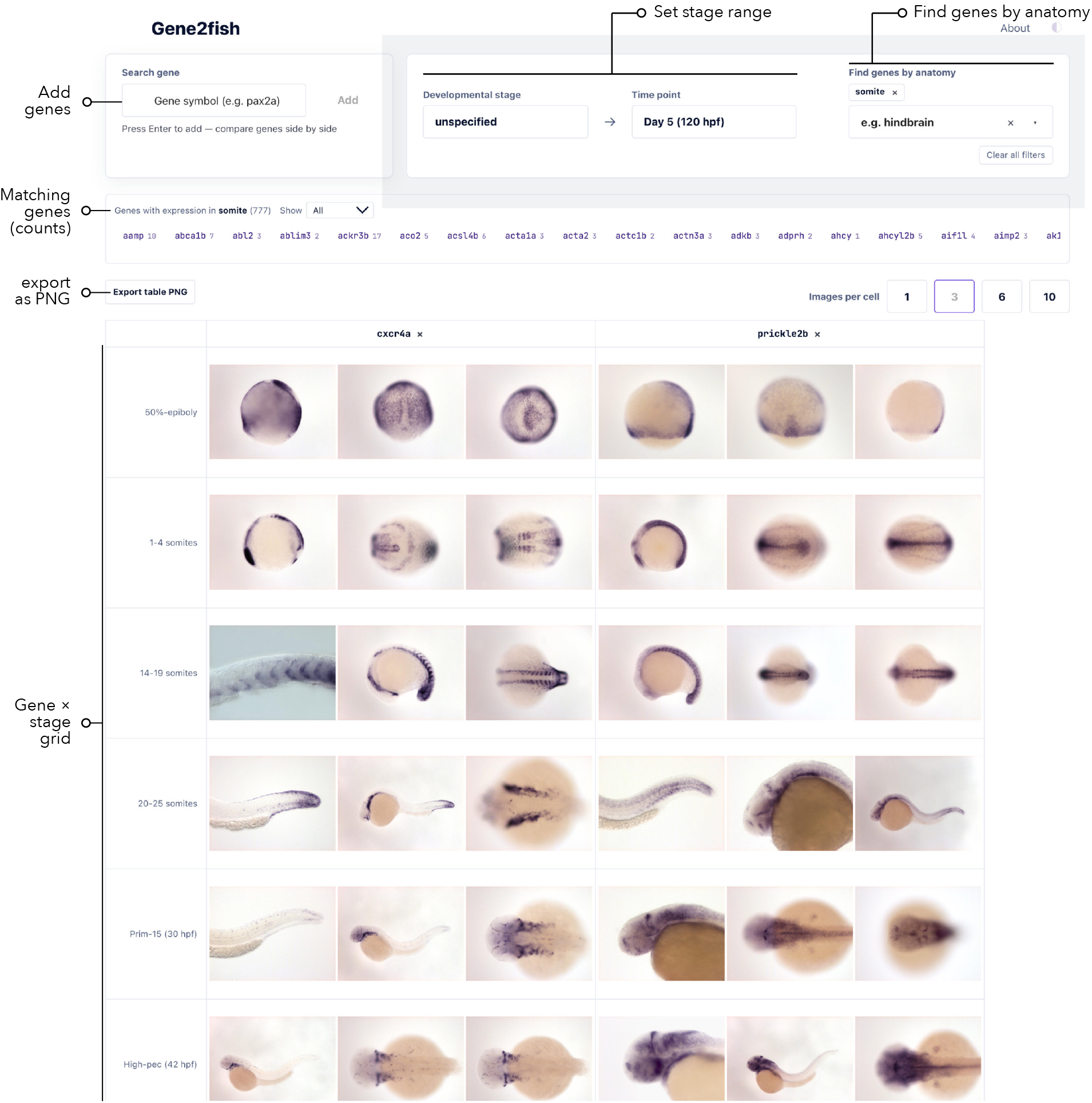
Main interface for gene2fish. The gene2fish web viewer of in situ hybridization data from the Thisse datasets. Users can examine gene expression across developmental stages or discover genes that are annotated for a selected anatomy term. Labels indicate the main controls: adding genes to the comparison, setting the developmental stage range, finding genes by anatomy, the strip of matching genes (each with its number of available images), exporting the current table as a PNG image, and the resulting gene *×* stage grid. Shown: *cxcr4a* and *prickle2b* at three images per cell, with the anatomy filter set to “somite” (777 genes).

With gene2fish, users can query multiple gene symbols, for example *cxcr4a* and *prickle2b*, which are then displayed in a comparative grid. Genes are displayed across columns and the rows correspond to zebrafish developmental stages. Each cell displays a representative image drawn from the Thisse datasets, or an empty placeholder if no data exist (see Materials and Methods for how images are assigned to stages and arranged) (Fig. 1; Supplementary Video 1). An **images-per-cell toggle** (1, 3, 6, or 10 images) allows users to display multiple images per stage when multiple preparations are available. A **stage range filter** (dual drop-down selectors populated from the developmental stage list) restricts visible rows to a user-defined developmental window, for example to focus on somitogenesis or pharyngula stages. In addition, users can choose from 227 ZFIN anatomy terms^22^ (including the generic terms “unspecified” and “whole organism”) to discover genes annotated to selected structures, or genes that are shared between anatomical regions (Fig. 1, Supplementary Fig. 2, Supplementary Table 1). As in ZFIN’s own expression search, selecting a term also returns genes annotated to its substructures by default (for example, *hindbrain* for *brain*), and annotations that ZFIN records as “expression not found” are never counted as expression. The resulting table can be exported as a PNG image for saving and sharing.

Clicking any image opens a full-resolution lightbox modal (Fig. 2) displaying the image alongside structured metadata: gene name and ZFIN gene ID, developmental stage, image preparation type, ZFIN anatomical annotation terms, EST/probe identifier and quality score, fish line, publication ID, image ID, corresponding human ortholog symbol, associated human diseases (inferred through orthology), and UniProt ID.

**Fig. 2:**
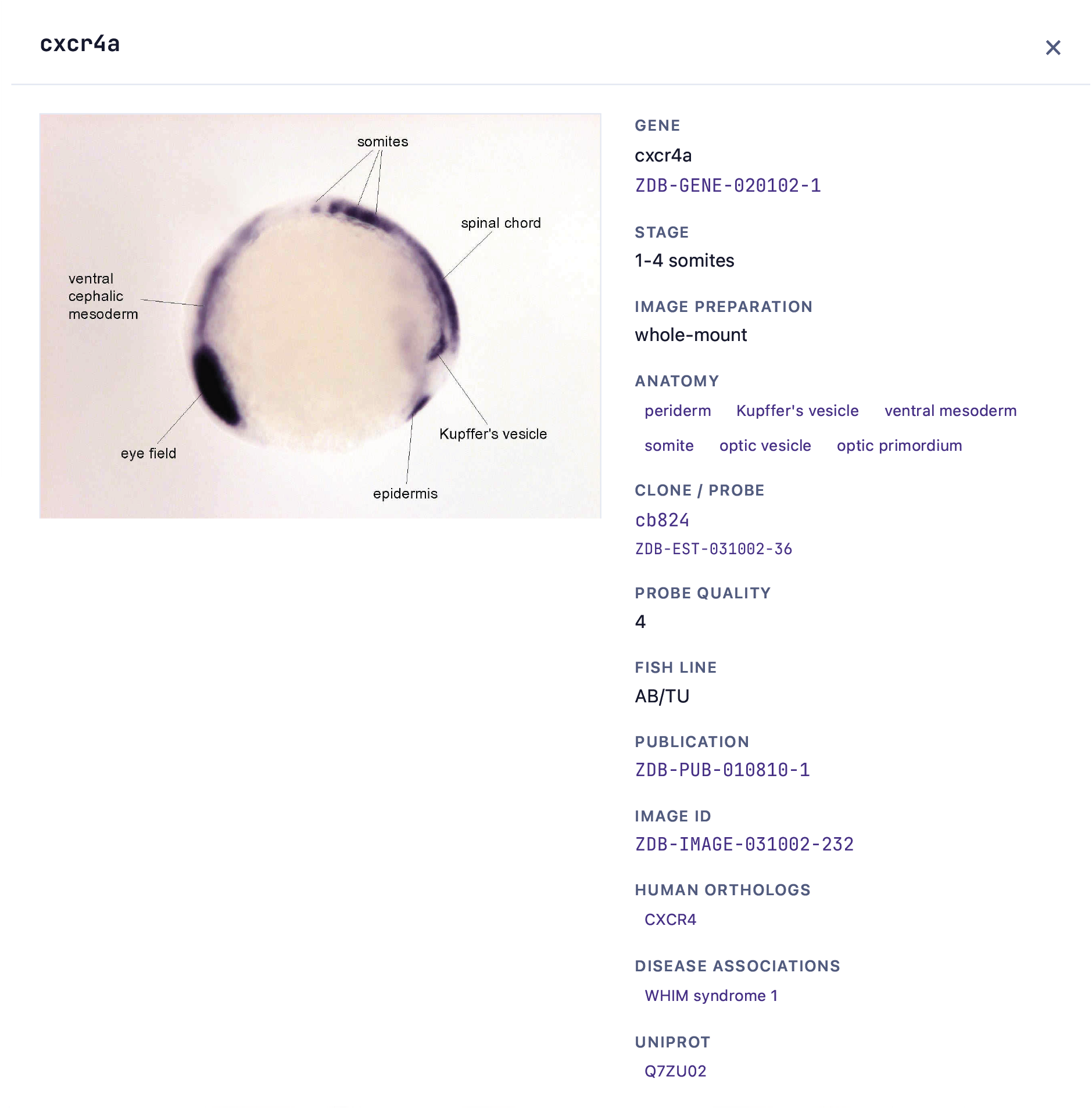
Image viewer. Clicking any image in the grid opens it at full resolution, here *cxcr4a* at the 1–4 somite stage, with the structure labels of the original Thisse *et al*. image where an annotated version exists. Alongside, the viewer lists the image’s ZFIN metadata: gene and ZFIN gene ID, developmental stage, preparation, the ZFA anatomy terms annotated for the figure (terms recorded as “expression not found” are listed separately), the clone or probe with its ZFIN quality rating (0–4), fish line, publication and image ID, and the human orthologs with their associated diseases (Disease Ontology terms inferred through orthology) and UniProt entries. Identifiers link to the corresponding records at ZFIN, NCBI and UniProt, and previous and next buttons step through all images shown in the grid.

For scRNA-seq workflows, a researcher validating a list of 10–20 cluster markers can survey their spatial patterns in a single browser tab rather than across 10–20 sequential ZFIN queries. Likewise, a researcher annotating anatomical structures can find the sets of genes annotated to them. Instead of visiting, or copying from, a separate ZFIN page for each gene, the user sees all genes at a glance and can immediately compare their expression domains across genes and tissues at matched developmental stages.

## Discussion

gene2fish addresses a routine bottleneck in relating zebrafish scRNA-seq data to spatial gene expression. Spatial datasets are becoming more common and will likely grow several-fold in the coming years, but they are currently restricted to early developmental stages and a limited number of genes. Until such data cover more stages and genes, gene2fish offers a practical alternative. The gene *×* stage grid aligns all genes along a shared axis of developmental time. This reveals changes in expression over time that single-stage lookups miss. The stage and anatomy filters then quickly narrow the view to the stages and structures most relevant to the experiment at hand.

ZFIN is widely used by zebrafish researchers, but its genecentric view can be a barrier for researchers from other disciplines. By showing the embryo images and their expression annotations together in one view, and using the detailed annotations of the Thisse datasets, gene2fish makes the exploration of gene expression across zebrafish tissues accessible to researchers outside the zebrafish field, including computational scientists.

Currently, gene2fish coverage is bounded by the Thisse high-throughput ISH collection; genes not assayed in that collection will yield empty cells, despite the presence of other RNA in situ hybridization datasets on ZFIN. Within the collection, coverage is uneven across stages, and a subset of images show pooled embryos of mixed stages annotated to an unspecified stage. Other data modalities, such as smFISH, RNAscope or spatial transcriptomics, are not yet integrated; future versions could include resources such as the MERFISHEYES atlas^13^. gene2fish is a browser and does not classify expression patterns automatically; interpreting them is left to the researcher.

gene2fish is open source and is available at https://zebrahub.sf.czbiohub.org/gene2fish with source code at https://github.com/czbiohub-sf/gene2fish. Images and annotations are provided by ZFIN^15^, are attributed to Thisse *et al*.^16–20^, and are used under the CC BY 4.0 license.

## Materials and Methods

### Data source

gene2fish uses the Thisse *et al*. high-throughput wholemount ISH collection at ZFIN^15–20^, that is, the images and annotations of five ZFIN direct data submissions (ZDBPUB-010810-1, ZDB-PUB-040907-1, ZDB-PUB-051025-1, ZDB-PUB-080220-1 and ZDB-PUB-080227-22). Metadata were taken from ZFIN’s public data reports (https://zfin.org/downloads), downloaded on April 14, 2026: the links between images and figures, the expression results shown in each figure, the developmental stage and anatomy of each expression result, and the expression details (gene, probe and probe quality, fish line, experimental conditions and publication), together with ZFIN’s stage, anatomy, gene, fish, condition, publication, human orthology, disease (via orthology), UniProt and gene alias reports. A Python script in the gene2fish repository (zfin image metadata extractor.py) joins these reports through their ZFIN identifiers into one record per image. The resulting index contains 53,595 images of 8,263 genes. All images show mRNA in situ hybridization on wild-type AB/TU embryos or larvae raised under standard conditions, and 99.6% are whole-mount preparations (the remainder being 166 sections, 14 live specimens and 30 unspecified preparations). This snapshot underlies all numbers reported here and is archived at Zenodo (doi:10.5281/zenodo.23378512). The deployed application rebuilds its index from the then-current ZFIN reports whenever it is redeployed, so counts in the live application may differ slightly as ZFIN updates its records.

### Developmental stages

ZFIN records a start and an end stage of the zebrafish staging series^21^ for each expression result. gene2fish assigns each image the earliest start stage among the expression results of its figure and maps it to the nearest of 37 canonical stages, from the 1-cell stage to adult; end stages are not used. Each of the figures in the collection covers a single stage, so all views in a figure fall into the same row of the grid. The grid shows only stages for which images exist, which for this collection are twelve stages from sphere to day 5 plus an “unspecified” row. Images whose start stage is recorded as the 1-cell stage are displayed as “unspecified”: in the Thisse submissions these are mostly images from the high-throughput screen^17^ showing pooled embryos of mixed stages (4,909 images), rather than onecell embryos.

### Anatomy annotations and search

ZFIN annotates anatomy per figure, using terms of the zebrafish anatomy ontology (ZFA)^22^. A Thisse figure shows one gene at one stage in one or more views, so all images of a figure share its anatomy terms; for each image, gene2fish collects the superterms and subterms of all expression results in its figure. ZFIN also records negative results (“expression not found”). The 227 terms annotated in the collection are offered for selection. By default, selecting a term also returns genes annotated to any of its substructures, following the *is a* and *part of* relations of the ZFA ontology (ZFIN release of September 28, 2026; 5,389 relations), as in ZFIN’s own expression search; an option restricts matching to the exact term. When several terms are selected, only genes annotated to all of them are returned, each with its number of matching images.

### Image selection and display

Within each gene *×* stage cell, images are arranged by preparation (whole-mount first), then by their number of positive anatomy annotations, then by ZFIN image identifier (most recent first), and users can display the top 1, 3, 6 or 10 images. The image viewer shows the metadata listed in the Results, including ZFIN’s probe quality rating on a scale from 0 (“probe is difficult to use”) to 4 (“simple to use, intense expression pattern restricted to a few structures”), which is not recorded for 30% of images. The complete view (genes, stage range, anatomy terms and images per cell) is encoded in the page URL, so it can be bookmarked and shared, and the current table can be exported as a PNG image that includes the attribution.

### Image hosting and attribution

ZFIN provided the Thisse image collection and permission to host it. To make image delivery reliable, the images are served unmodified from a Biohub cloud storage bucket that contains only this collection; an image absent from the bucket is loaded directly from ZFIN. The grid displays ZFIN’s medium-resolution version of each image and the image viewer the full-resolution original. Every image links to its ZFIN record, and the application and exported tables include the attribution wording agreed with ZFIN. Images and annotations are used under the CC BY 4.0 license.

### Software implementation and deployment

The backend is written in Python (version 3.11) with FastAPI; it loads the index into memory at startup and serves gene, anatomy, stage and image queries through a REST API. The frontend is a React (version 18) single-page application built with Vite. Both are packaged in one container image, and the metadata index is built when the container image is built, so updating the data requires only a rebuild. The application runs on Biohub’s cloud infrastructure and is embedded in the Zebrahub portal. The repository includes backend unit tests (pytest) and browser end-to-end tests (Playwright). The version described here (v0.14.2) is archived at Zenodo (doi:10.5281/zenodo.23380713).

### Dataset statistics

The numbers of images and genes per stage (Supplementary Fig. 1) and the numbers of genes per anatomy term (Supplementary Fig. 2 and Supplementary Table 1) were computed from the April 14, 2026 snapshot with Python scripts (using matplotlib) that are deposited with the dataset at Zenodo (doi:10.5281/zenodo.23378512). Counts per anatomy term exclude “expression not found” annotations, and the substructure counts in Supplementary Table 1 use the ontology relations described above.

### Use of AI tools

Generative AI tools (Claude, Anthropic) assisted with editing the manuscript text and Supplementary Video 1, and with the design and development of the gene2fish software. All AI-assisted content was reviewed and verified by the authors, who take full responsibility for this work.

## Supporting information

Supplementary Video 1. Walkthrough of the gene2fish interface.

## Data and resource availability

The gene2fish web application is available at https://zebrahub.sf.czbiohub.org/gene2fish. Source code is available at https://github.com/czbiohub-sf/gene2fish under the MIT license; the version described here (v0.14.2) is archived at Zenodo (doi:10.5281/zenodo.23380713).

All images and annotations are from public ZFIN data reports (downloaded April 14, 2026). The metadata index used in this study, together with the anatomy hierarchy, Supplementary Table 1 and the scripts used for the supplementary figures and table, is available at Zenodo (doi:10.5281/zenodo.23378512) under the CC BY 4.0 license.

## Acknowledgements

We thank the ZFIN team, who provided the image collection, technical support and feedback on our application: Yvonne Bradford, Ryan Martin, Ryan Taylor, Christian Maass, and Monte Westerfield. We are also immensely grateful to the original authors who created the in situ images: C. Thisse, B. Thisse, S. Pflumio, M. Fürthauer, B. Loppin, V. Heyer, A. Degrave, R. Woehl, A. Lux, T. Steffan, X. Q. Charbonnier, and G. J. Wright. We thank S. Gordon Leary, Reinier John Grona, Jia Le Lim and Allyson Quinn Ryan for feedback on gene2fish. We thank the Biohub and its donors, Priscilla Chan and Mark Zuckerberg, for funding this work.

## Author contributions

V.J. conceived the research and developed the prototype of gene2fish. W.D.F.R. and L.L. engineered the application. K.A. and B.I. published gene2fish on Zebrahub and maintain it. V.J. wrote the original draft of the paper. V.J., L.A.R., W.D.F.R. and L.L. reviewed and edited the manuscript. V.J. led the project. L.A.R. supervised the research. R.G.-S. supervised the software engineering. Data were curated and shared by ZFIN.

## Supplementary Figures

**Supplementary Fig. 1:**
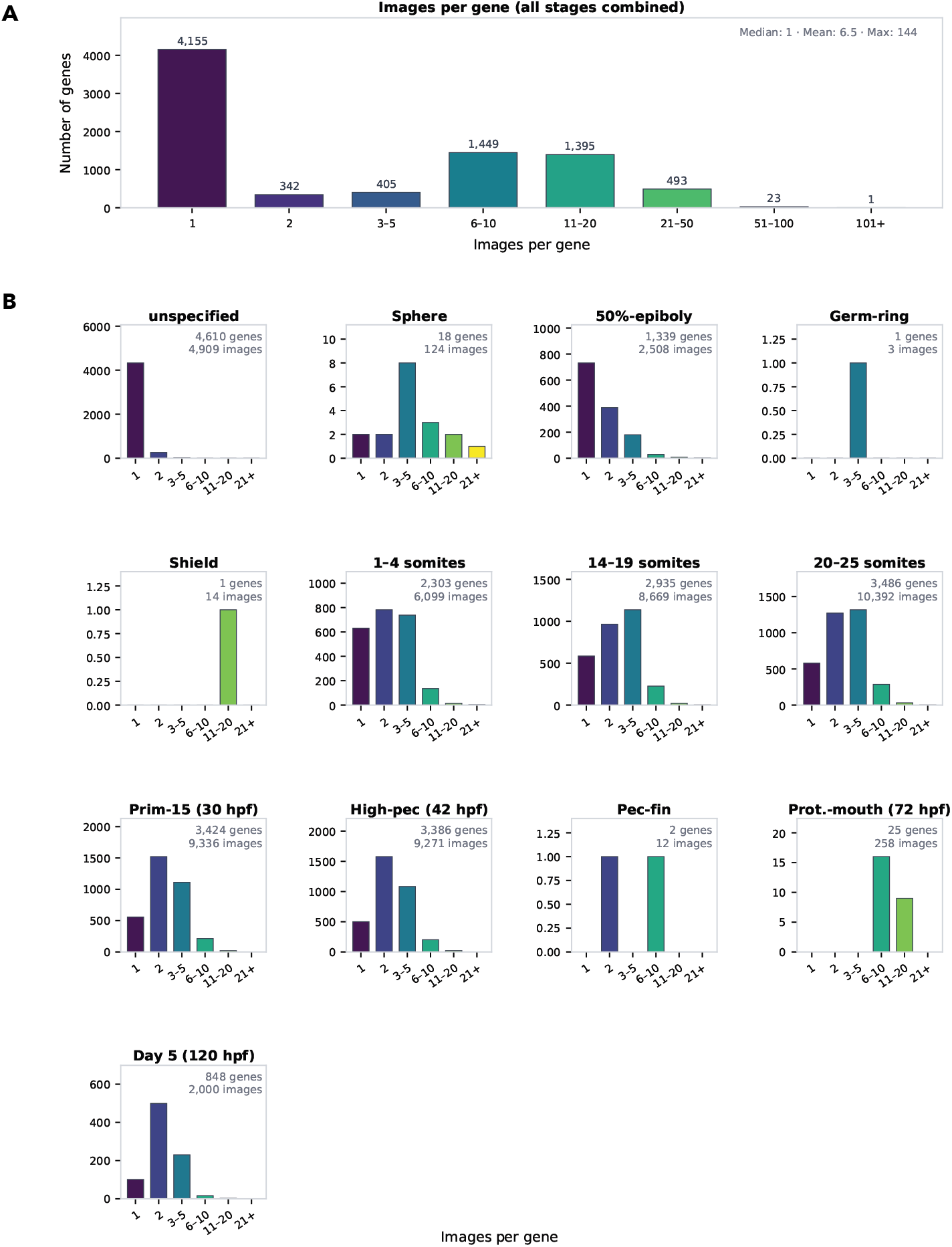
Image coverage of the Thisse ISH collection per gene and across developmental stages. (**A**) Number of images per gene, all stages combined, across the full dataset (8,263 genes, 53,595 images). The majority of genes are represented by a single image (median 1, mean 6.5, maximum 144 images per gene), while a subset of well-characterized genes is covered by 6–20 images across multiple preparations. (**B**) The same distribution for each developmental stage, in bins of 1, 2, 3–5, 6–10, 11–20 and more than 20 images per gene; each panel plots the number of genes against images per gene and gives the numbers of genes and images at that stage (top right). Coverage is highest at somitogenesis and pharyngula stages (1–4 somites through High-pec). “Unspecified” denotes mostly images from the high-throughput screen^17^ showing pooled embryos of mixed stages.

**Supplementary Fig. 2:**
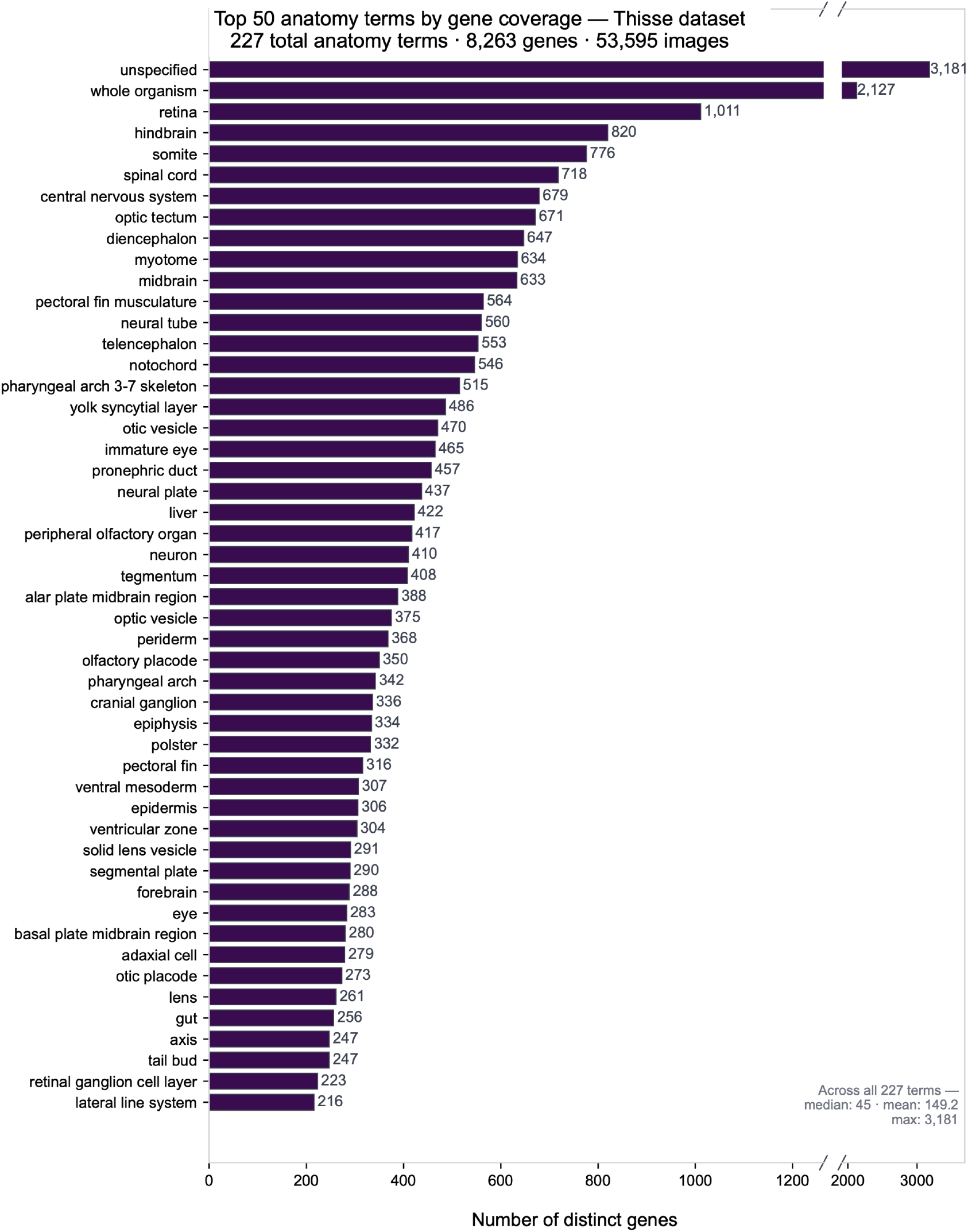
Dataset coverage of the Thisse ISH collection across ZFIN anatomy terms. The 50 ZFIN anatomy terms with the most genes, ranked by the number of distinct genes with at least one image annotated to the term, out of the 227 terms annotated across the dataset (8,263 genes, 53,595 images); annotations recorded by ZFIN as “expression not found” are excluded. The x axis is broken to accommodate the two generic terms “unspecified” and “whole organism”. Across all 227 terms, the median is 45 genes per term (mean 149.2, maximum 3,181). The full ranked list of all 227 terms, including counts with substructures, is provided in Supplementary Table 1.

## Supplementary Videos

**Supplementary Video 1.**
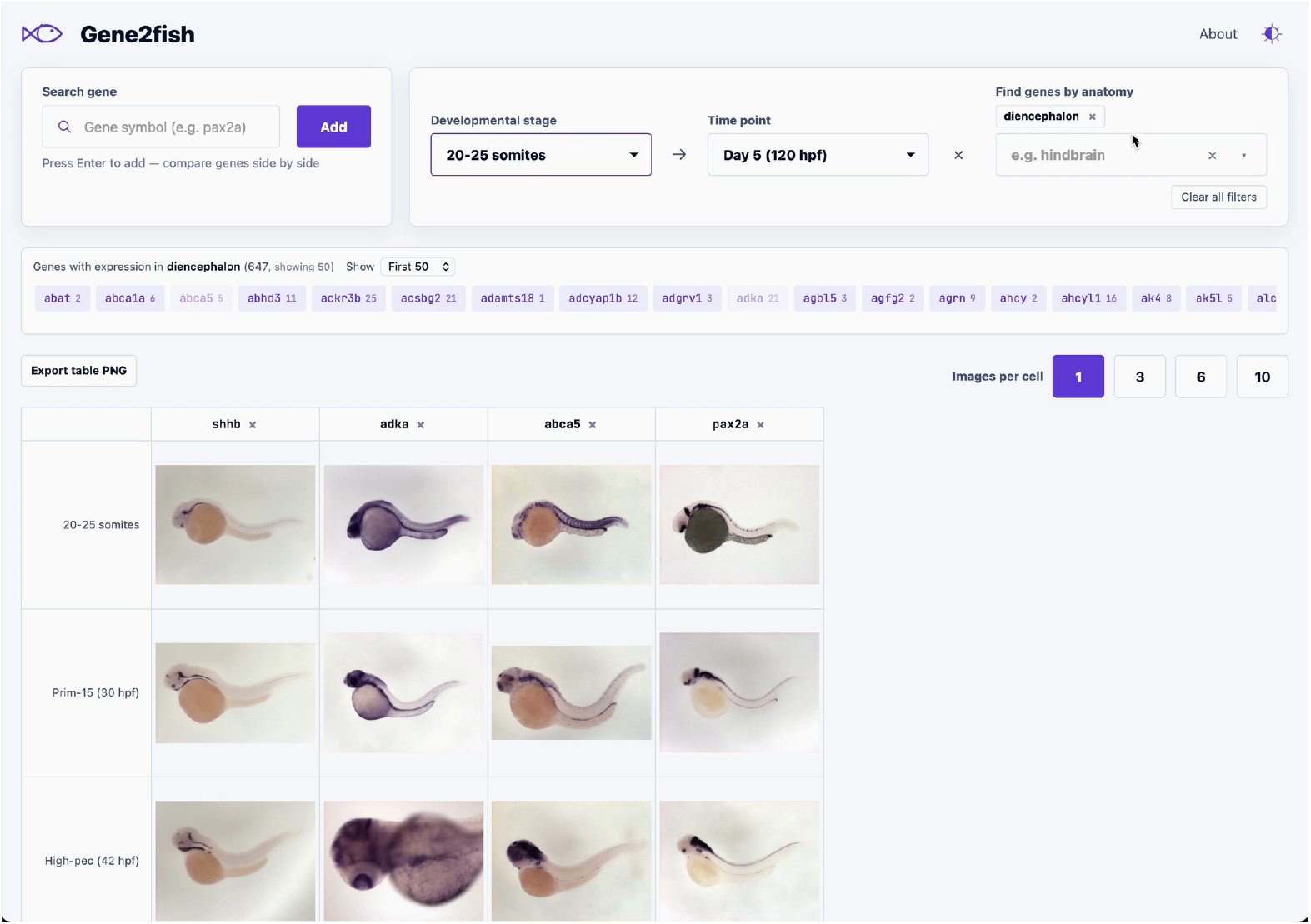
Walkthrough of the gene2fish interface. Starting from the landing page, the example gene *shhb* is loaded and its Thisse ISH images are displayed across developmental stages; hovering over an image shows its stage, ZFIN image ID and publication ID. The images-per-cell toggle is then increased to display multiple preparations per stage. Selecting the anatomy term “diencephalon” lists the genes annotated to that structure (647 genes; first 50 shown, with the number of images per gene), from which *adka* and *abca5* are added to the grid with a single click; *pax2a* is added through the gene search field with autocompletion. The developmental stage range is then restricted to 20–25 somites through Day 5 (120 hpf), and clicking an image opens the lightbox with its structured ZFIN metadata. Screen recording of https://zebrahub.sf.czbiohub.org/gene2fish; 78 s, silent. The frame shown above is at 63 s. File: Supplementary Video 1 gene2fish.mp4.

## Supplementary Data

**Supplementary Table 1.**
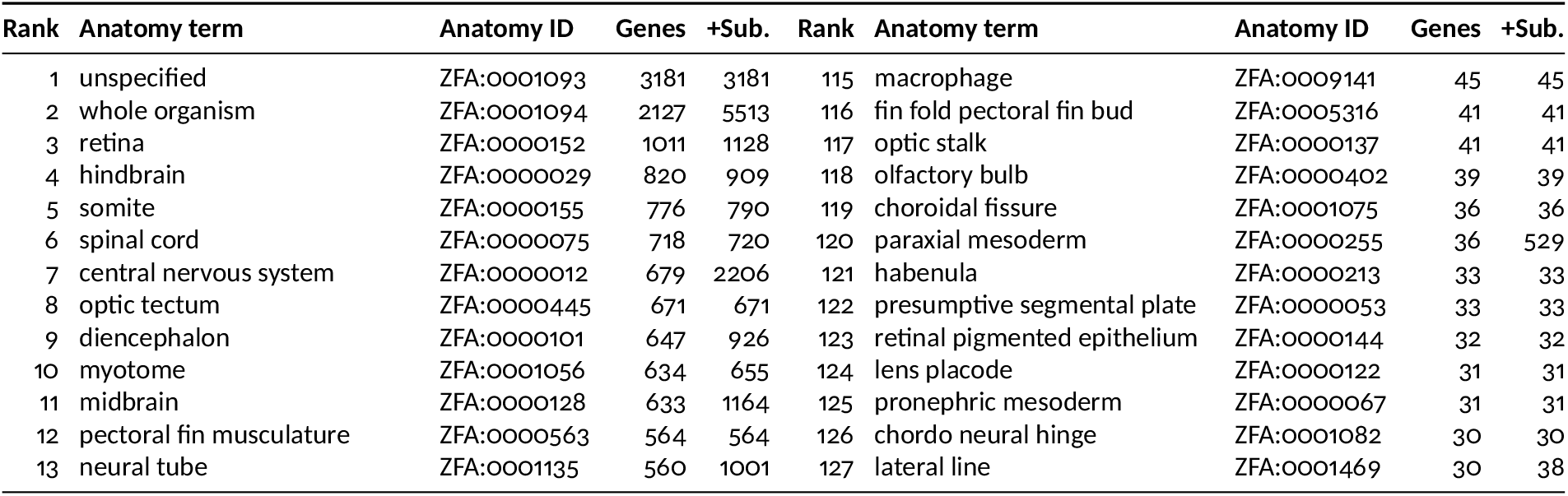
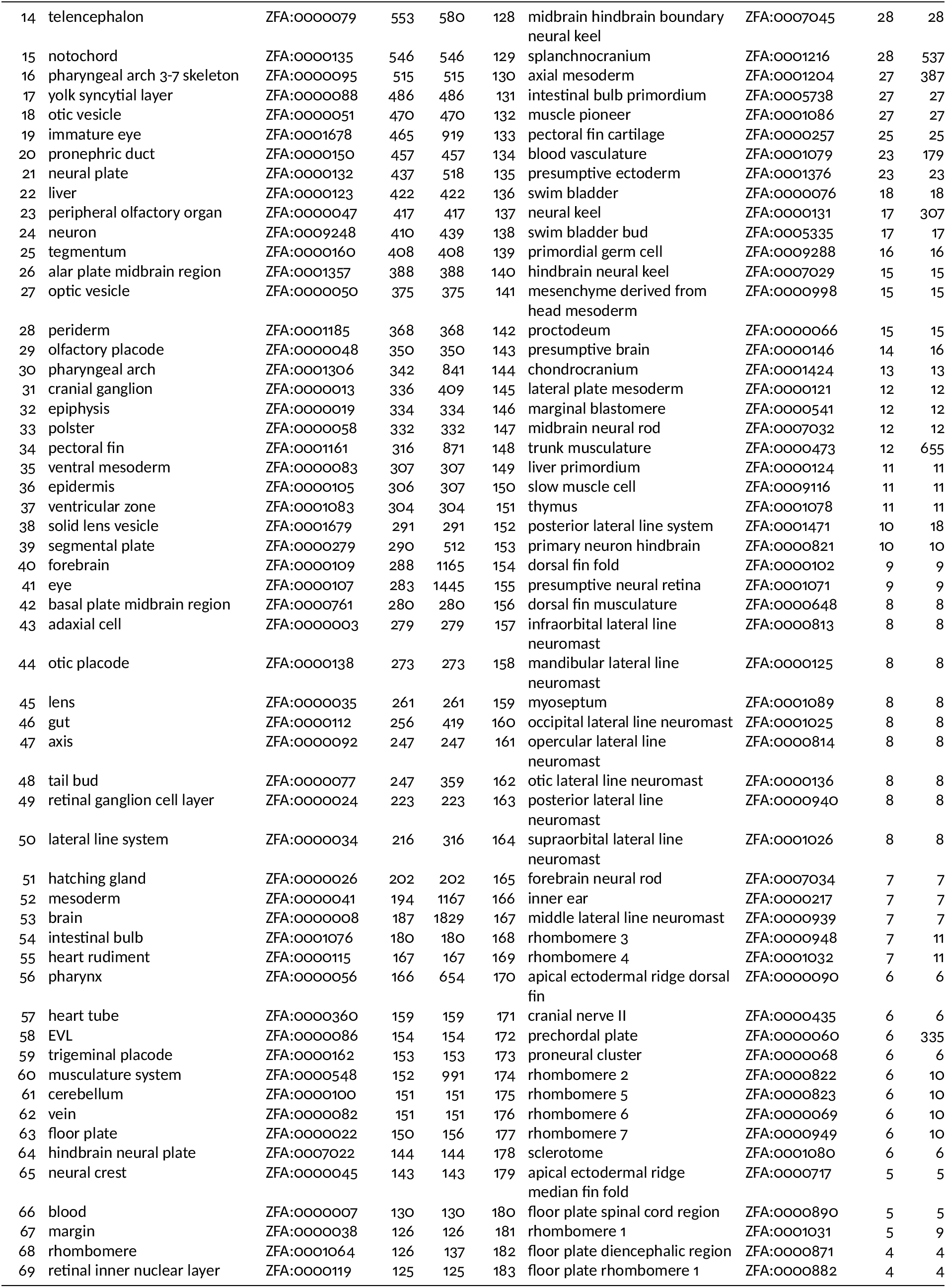
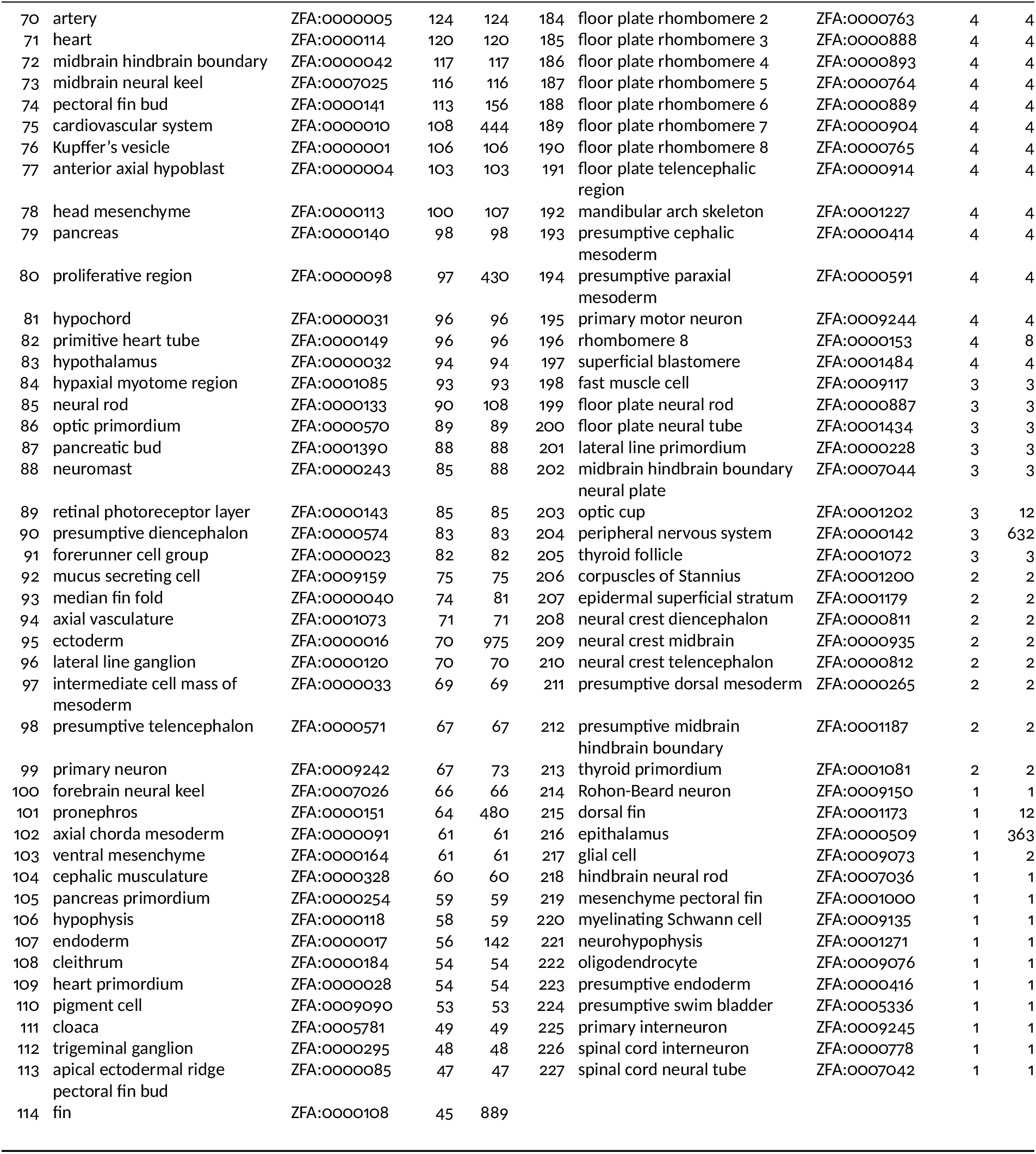
All 227 ZFIN anatomy terms browsable in gene2fish, ranked by the number of genes with at least one image annotated to that term (**Genes**). **+Sub**. counts genes annotated to the term or to any of its substructures (ZFA *is a*/*part of* relations), which is what gene2fish returns by default. Annotations recorded by ZFIN as “expression not found” are excluded.

